# Stability and Control of Biomolecular Circuits through Structure

**DOI:** 10.1101/2020.11.04.368381

**Authors:** Fangzhou Xiao, Mustafa Khammash, John C. Doyle

## Abstract

Due to omnipresent uncertainties and environmental disturbances, natural and engineered biological organisms face the challenging control problem of achieving robust performance using unreliable parts. The key to overcoming this challenge rests in identifying structures of biomolecular circuits that are largely invariant despite uncertainties, and building feedback control through such structures. In this work, we develop the tool of log derivatives to capture structures in how the production and degradation rates of molecules depend on concentrations of reactants. We show that log derivatives could establish stability of fixed points based on structure, despite large variations in rates and functional forms of models. Furthermore, we demonstrate how control objectives, such as robust perfect adaptation (i.e. step disturbance rejection), could be implemented through the structures captured. Due to the method’s simplicity, structural properties for analysis and design of biomolecular circuits can often be determined by a glance at the equations.

## I. Introduction

Both natural and engineered cells face the challenge of achieving robust performance using unreliable parts [1]–[3]. In particular, the regulatory biomolecular circuits used in a cell to achieve desired behavior face large parameter uncertainties due to environmental disturbances, and unknown or unintended interactions with background cell circuits.

Although feedback control has been successfully applied in electrical and mechanical engineering to achieve robust performance [4], it faces a new challenge in biological engineering that the parts are highly unreliable. Therefore it is essential to identify key structures of the uncertain behaviors in biomolecular circuits, so that feedback control can be built on top of them.

Previous studies have identified several important structures of biomolecular circuits. While reaction rates tend to vary due to environmental disturbances, the stoichiometry of reactions are invariant, since they are results of atomic compositions of molecules. Indeed, stoichiometry could be robustly identified from experimental data and is often considered as structural information of a chemical reaction network [5], [6]. In addition, although rate constants and reactant concentrations vary due to disturbances, they often can be reliably determined or controlled up to orders-of-magnitude [7]. Lastly, the many reactions that happen in a cell often happen at different time scales, making descriptions of a circuit’s behavior amenable to time-scale separation [8]. The most robust separation of time scales is the one between binding reactions and catalysis reactions, as exemplified by the Michaelis-Menten approximation, which has served as the foundation of dynamic modeling of biochemical reactions for over 100 years [9]–[11]. A simple physical argument is that binding reactions are fast as they only involve low-energy interactions such as hydrogen bonds, while catalysis changes high-energy covalent bonds, therefore slower.

The structures mentioned above need to be synergistically integrated in a cohesive mathematical framework in order to analyze or design robustly performing biomolecular circuits using unreliable parts. In particular, it needs to connect structures with dynamical properties of the system. This difficult challenge, yet to be overcome, is the central cause for a major gap between the mathematical languages theorists use, and the mental pictures and diagrams that experimentalists use to guide their circuit designs and implementations [12]–[14].

This work provides initial results that could serve as a first attempt at filling this gap. In particular, we aim at building mathematical concepts that are tailored for these quintessentially biological structures.

In the following, we define a general class of systems, named birth-death systems, that emphasize the production and degradation of biomolecules, in Section II-A. In Section II-B, we use log derivatives to capture the structure in production and degradation fluxes’ dependence on reactant concentrations. In Section III, we show how log derivatives relate to a strong notion of stability of fixed points. Lastly, in Section IV, we show how feedback control could be built on top of structures to achieve control goals such as robust perfect adaptation, biologists’ term for step disturbance rejection.

A companion work with a focus on studies of examples that cater to a more biological audience is [15].

## II. Structure in Biomolecular Systems

We begin by introducing the definition of birth death systems. We do so through chemical reaction networks (CRNs) [16] to have explicit biological interpretations, although birth-death systems do not rely on the formalism of CRNs.

### A. Chemical Reaction Networks and Birth Death Systems

A CRN is a collection of reactions of the form

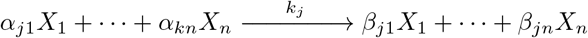

where *X*_*i*_, *i* = 1*,…, n* denote chemical species, *j* = 1*,…, m* index reactions, 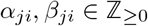 denote the number of *X*_*i*_ molecules needed as reactant or produced as product in reaction *j*, and *k*_*j*_ ∈ ℝ_>0_ is reaction rate constant of of reaction *j*. We denote ***α***_*j*_ = *α_j_*_1_ · · · *α_jn_* as the reactant stoichiometry vector for reaction *j*, and similarly define ***β***_*j*_ for product Vector. We define ***γ***_*j*_ = ***β***_*j*_ − ***α***_*j*_ as the stoichiometric vector of reaction *j*, and 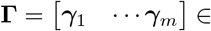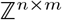 is the stoichiometric matrix.

The deterministic rate equation of the CRN is

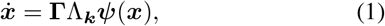

where *x*_*i*_ ∈ ℝ_≥0_ is the concentration of species *X*_*i*_, Λ_***k***_ := diag {***k***} is a diagonal matrix with reaction rate constants *k_j_* as entries, and 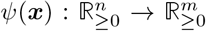 denote how the rate of reactions depend on concentrations.

A commonly used specification for *ψ_j_* (***x***) is the law of mass-action, which is applicable to a wide range of scenarios [17]. It says *ψ_j_*(***x***) = ***x***^***α**j*^, where we denote 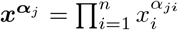

Since concentrations of biomolecules change by production and degradation reactions, we could re-write the dynamics as follows:

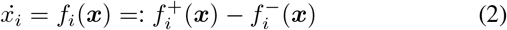

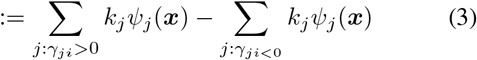

where we have collected terms from reactions producing *X*_*i*_ into 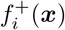 and terms from reactions degrading *x*_*i*_ into 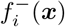.

The physical interpretation of the variables *x*_*i*_ as concentrations dictate that they remain non-negative, therefore the positive orthant is forward invariant. A necessary and sufficient condition is *f_i_*(***x***) ≥ 0 whenever *x*_*i*_ = 0. It is also natural to assume that each species has at least one production reaction and at least one degradation reaction. This yields the following definition for birth-death systems.

#### Definition 1

A birth-death system is a dynamical system of the form (2) where the production and degradation rates 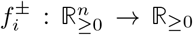 are analytic and globally non-negative, and *f*_*i*_ (***x***) 0 whenever *x*_*i*_ = 0.

The definition of a birth-death system emphasizes the structure that each concentration variable is regulated by two processes, production and degradation. Understanding the dynamics of a birth-death system then comes down to characterizing how production and degradation rates 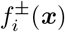 depend on the concentrations ***x***. In the following section, we use a simple example to illustrate that the dependence of 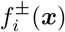 on ***x*** is highly structured, and this structure could be formalized through log derivatives.

### B. Log derivatives formalize regimes of regulation under time-scale separation

Production and degradation of molecules happen through enzymatic catalysis [10]. In the following, we consider the simplest regulation of enzymatic catalysis to illustrate the structure in 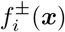’s dependence on ***x***.

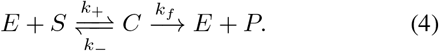

Here *E* is the enzyme, *S* is substrate, *C* is the complex formed from *E* and *S* binding together, and *P* is the product molecule formed.

To proceed, we use time-scale separation that binding reactions tend to be much faster than catalysis reactions. This entails the following equations:

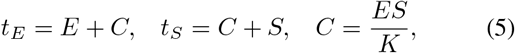

where *t_E_* is the total concentration of enzyme *E*, *t_S_* is total concentration of substrate *S* in free or bound forms, and *K* is the dissociation constant *K_d_* or its variants such as *K_M_*, based on details of which part of *C* dynamics is fast [18].

To connect with the notation of birth-death systems, we denote *x_P_* as the concentration of *P*, *x_E_* = *t_E_* as the total concentration of *E*, and *x_S_* = *t_S_* as the total concentration of *S*. Then *x_P_*’s dynamics is

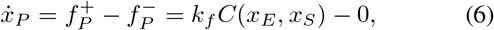

where *C*(*x_E_, x_S_*) is the steady state expression determined by Eq (5).

In order to understand how 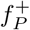, the production rate of *x_P_*, depends on *x_E_* and *x_S_*, we need to solve for *C* in terms of *t_E_* and *t_S_* in Eq (5). A classical way to approach this is the Michaelis-Menten approximation [18], which assumes the total concentration of the substrate is much higher than that of the enzyme, i.e. *t_S_ ⨠ t_E_*. This implies *t_S_ ≈ S*, therefore Eq (5) solves to be

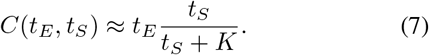

This expression could be intuitively understood as containing two regimes. One has *t*_*S*_ ≫ *K*, so that *C* ≈ *t*_*E*_. This is constant in *t_S_*, therefore “substrate-saturated”. The other one has *t_S_* ≪ *K*, so that 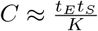. This is linear in *t_S_*, therefore “substrate-sensitive”. We note that these two regimes have distinct exponents in *t_E_* and *t_S_* : (1, 0) for the saturated regime, and (1, 1) for the sensitive regime (see Figure 1).

**Fig. 1:**
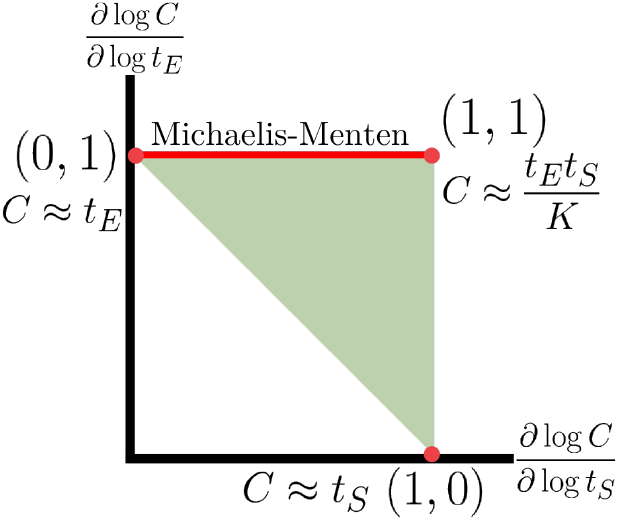
The log-derivative polytope of the complex *C* with respect to *t*_*E*_ and *t*_*S*_ defined by steady state equations in Eq (5). A point in this space represents the sensitivity of the steady-state *C* concentration to changes in the total concentration of *E* or *S*. The green triangle marks the possible sensitivity values the system can admit. The edges of the triangle represent different assumptions about the saturation of the species. The edge marked by the red line is the range of log derivatives covered by the Michaelis-Menten formula. Red dots mark the vertices. The expressions next to the vertices correspond to the three regimes.

Therefore, although 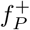, the production rate of *x_P_*, depend on concentrations and rates that tend to be uncertain in Eq (7), the fact that there are two regimes, each with distinct exponents in the concentrations, is structural. Indeed, the exponents fundamentally come from the stoichiometry of the binding reaction in Eq (4). In addition, the condition such that one regime is valid, such as *t*_*S*_ ≫ *t*_*E*_, *K*, only depend on the orders of magnitude of the concentrations and rates, therefore could be reliably determined or controlled.

Now, we would like to describe these regimes and their exponents in a formal way. For this purpose, we introduce log derivatives as differential analogues of exponents. For example, from Eq (7) we calculate

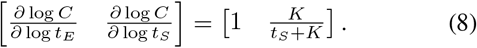

When *t*_*S*_ ≫ *K*, we obtain log derivative (1, 0); while when *t*_*S*_ ≪ *K*, we obtain (1, 1). So log derivatives exactly capture the exponents of the regimes in a continuous way.

With the tool of log derivatives in mind, we could actually go back and obtain more general results than the Michaelis-Menten approximation. Indeed, due to the assumption that *t*_*S*_ ≫ *t*_*E*_, we missed the third “enzyme-saturated” regime: when *t*_*E*_ ≫ *t*_*S*_, *K*, we have *C* ≈ *t*_*S*_. Capturing this regime is important if the *t*_*S*_ ≫ *t*_*E*_ assumption does not hold all the time, such as when *S* and *E* are molecules of similar abundance in protein binding, or when the cellular circuit has highly dynamic behavior during nutrient shifts or shock responses [19], [20].

To capture all three regimes, we need to do away with assumptions like Michaelis-Menten. Although the steady state equations in Eq (5) can be directly solved in this simple case, this procedure results in a cumbersome expression that does not generalize to more complicated cases. More importantly, the explicit solution obtained hides the structures in the exponents of the regimes mentioned above. In contrast, the differential description through log derivatives can capture all three regimes while describing the exponents. Indeed, applying implicit function theorem to Eq (5) to solve for the expression of *E, S, C* in terms of *t*_*E*_, *t*_*S*_, *K* yields the following result:

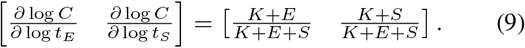

This shows that the log derivatives of *C* with respect to *t_E_* and *t_S_* take values inside a triangle (see Figure 1), and the exact location in the triangle depends on the particular values of *t_E_, t_S_* and *K*. Completely in accordance with our intuition, the vertices of this triangle correspond to the three regimes described earlier. In particular, we see that the Michaelis-Menten approximation is just one edge of this triangle, a strict subset of the behaviors captured by the log derivatives.

The fact that the log derivatives form a triangle, i.e. the set of convex combinations of the three vertices, suggests that the full behavior of the enzyme regulation could be seen as combinations of the three regimes corresponding to the three vertices. Indeed, when the corresponding asymptotic conditions are satisfied, the behavior of the enzyme regulation is essentially the same as the simple monomials at the vertices. Extending this to all production and degradation fluxes, we see that a general birth-death system could be seen as having several regimes, each corresponding to a simple system with constant exponents. Depending on the location of the state, the system could be approximated by one or another simple system corresponding to its closest regime. Hence the following definition.

#### Definition 2

A *simple birth-death system* is a birth-death system with 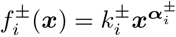, where 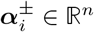 is a constant where vector, and 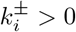 is a positive constant.

Simple birth-death systems have the advantage that their log derivatives can be directly read off from the exponent vector in the rate functions. In contrast, obtaining the set of log derivatives that emerges directly from binding networks is nontrivial in general. Next, we show that log derivatives do form easily-identified polytopes in most models of biological circuits, where Hill functions from Michaelis-Menten approximations are used.

### C. Basic facts about log derivatives

Here are some basic calculations to facilitate intuition about log derivatives.

If 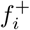 is a monomial, i.e. 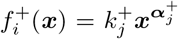, then 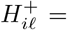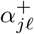 for *ℓ* = 1,…, *n*. In other words, the log derivative vector 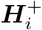 for the production of *X*_*i*_ is the exponent vector 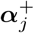 independent of the rate constant 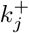 or concentration ***x***. This case corresponds to simple birth death system. Physically, this case could happen when *X*_*i*_ has only one production reaction. Then 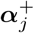 is the reactant stoichiometry vector for that production reaction.

If 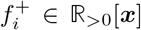 is a multivariate polynomial in with *x* positive coefficients, i.e. 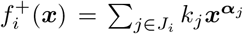 for index set 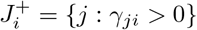 of all reactions producing *X*_*i*_, then

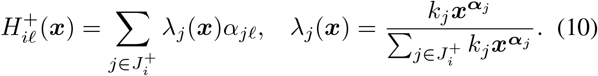

Since *λ*_*j*_ > 0 and sums to one, the log derivative vector for the production rate of *X*_*i*_ is the convex combination of the reactant vectors of all *X*_*i*_-producing reactions. In other words, 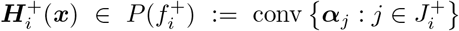, where 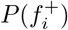 is the Newton polytope of polynomial 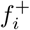. Although the location in the polytope depends on *k_j_* and ***x***, the polytope itself depends on reactant vectors ***α**_j_* alone. We note that newton polytopes are fruitful tools in analysis and optimization of polynomial equations, dynamical systems, and CRNs [21]–[23].

If 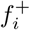 is a rational function with the numerator as one term of the denominator polynomial, i.e.

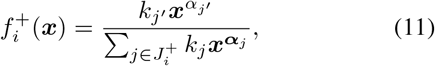

which typically arises from time-scale separations and Michaelis-Menten type approximations, then

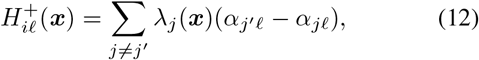

where *λ_j_* is the same as before. In this case, the log derivative vector for production of *X*_*i*_ is the convex combination of all reactions’ reactant v tors minus the numerator reactant vector: 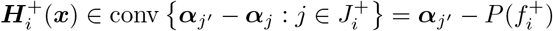.

The above calculations enable writing down log derivatives immediately after a glance at the equation in many cases, making log derivatives easy to use.

## III. Structure and Fixed Point Stability

We have shown that the structures of biomolecular circuits could be elegantly captured via log derivatives. In the following, we discuss how log derivatives connect with the stability of fixed points in birth-death systems. In particular, we show that log derivatives could certify structural stability of a fixed point: stability that is independent of concentrations and rates.

### A. Linearization and logarithmic derivatives

We first assume that the birth-death system has a positive fixed point 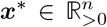 such that ***f*** (***x***^∗^) = 0, with positive production and degradation fluxes: 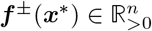

Since the stability of a fixed point is determined by the eigenvalues of the linearized dynamics at that fixed point, we express the linearization of a birth-death system in terms of log derivatives. We introduce the log derivative map

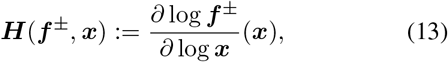

where log is applied component-wise. The log derivative map takes a positive function ***f*** ^±^ and a point ***x*** in its domain to the function’s log derivative at this point. For simplicity, we denote ***H***^+^ := ***H***(***f*** ^+^*, **x***), ***H***^−^ := ***H***(***f*** ^−^*, **x***), and ***H*** := ***H***^+^ − ***H***^−^.

We calculate that derivatives could be expressed in terms of log derivatives as

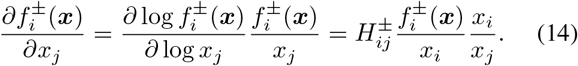

In matrix form, this is

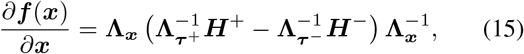

where 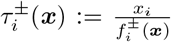 are time-scales of *X*_*i*_’s production or degradation fluxes [24].

At a fixed point ***x***^∗^, we have ***f*** ^+^(***x***^∗^) = ***f*** ^−^(***x***^∗^), so we could define ***τ*** := ***τ*** ^+^(***x***^∗^) = ***τ*** ^−^(***x***^∗^) as the vector of steady-state time-scales at fixed point ***x***^∗^. Therefore, we have

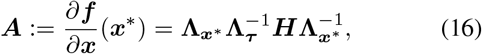

relating linearized dynamics ***A*** to log derivative matrix ***H***.

Let us define 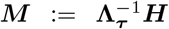. Since ***A*** and ***M*** are similar to each other, they have the same eigenvalues. So we immediately see that the fixed point ***x***^∗^ is stable if and only if ***M*** is Hurwitz. Therefore, the stability of the fixed point ***x***^∗^ depends only on ***M***, which is nicely split into two parts: the time-scales in 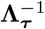, and the log derivatives in ***H***. Since the time-scales come from uncertain rates, while log derivatives capture reliable structures of the system, this prompts the following definition:

#### Definition 3

A fixed point ***x***^∗^ of a birth-death system is *structurally stable* if it is stable for all positive analytic rate functions ***f*** ^±^ that leave the log derivative matrix ***H*** at ***x***^∗^ invariant.

As the variations in rates could only change time-scales ***τ*** in matrix ***M***, the definition of structural stability exactly corresponds to that the log derivative matrix ***H*** is (multiplicative) *D*-stable in matrix analysis. In other words, left multiplication of ***H*** by arbitrary positive diagonal matrices results in a Hurwitz matrix (see extensive review by [25]). *D*-stability has been extensively studied since the very beginning of control theory, yet a clean necessary and sufficient characterization has not been found. This is in part due to the topological pathology of this property, that the set of *D*-stable matrices is neither closed nor open. A good sufficient condition to *D*-stability that characterizes a topologically nice (open) set of matrices is diagonal stability:

#### Definition 4

A matrix ***H*** is *diagonally stable* if there exists a positive diagonal matrix ***P*** such that

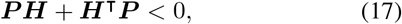

where < 0 for matrices denote negative definiteness.

Since Eq (17) is a linear matrix inequality, scalable numerical solution algorithms are available off-the-shelf.

A theorem summarizes above discussions.

#### Theorem 5

***H*** is diagonally stable implies the fixed point ***x***^∗^ is structurally stable. ***H*** is diagonally stable if and only if ***A*** is diagonally stable.

*Proof:* We calculate that

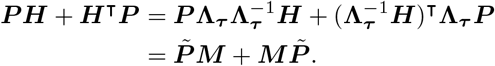

Therefore, ***H*** is diagonally stable is equivalent to ***M*** is diagonally stable, which is equivalent to ***A*** is diagonally stable as they are similar through positive diagonal matrix multiplications. Also, this calculation shows ***H*** is diagonally stable implies that for any positive diagonal matrix 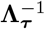, ***M*** satisfies 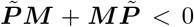 for 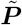 that is positive definite, therefore ***M*** is Hurwitz by basic Lyapunov theory.

There are a few special cases of the above theorem that is worth mentioning due to their simplicity.

#### Corollary 6

Any of the following conditions imply ***x***^∗^ is structurally stable.

1. ***H*** is triangular with negative diagonal entries.
2. ***H*** is symmetric and negative definite.
3. The symmetrization of ***H***, 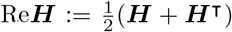, is negative definite.

Structural methods could be easily extended to certify fixed points’ structural instability as well. This is because when ***H*** has no purely imaginary eigenvalues, any symmetric matrix ***P*** that satisfies Eq (17) has inertia (i.e. signs of eigenvalues’ real parts) that are opposite to the inertia of ***H***, as stated in the Ostroski-Schneider theorem in matrix analysis [25], [26]. Therefore, if there exists a diagonal matrix ***P*** with at least one negative entry satisfying Eq (17), then ***H*** is not Hurwitz, and 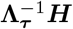 is not Hurwitz for all positive vectors ***τ***. We summarize this into the following theorem.

#### Theorem 7

Given a birth-death system as in (2) with log derivative matrix ***H*** at a positive fixed point ***x***^∗^, if ***H*** has no purely imaginary eigenvalues and there exists a diagonal matrix ***P*** with at least one negative entry such that Eq (17) holds, then ***x***^∗^ is structurally unstable, i.e. its linearized dynamics is unstable for all positive analytic functions ***f*** ^±^ that keeps ***H*** invariant at ***x***^∗^.

### B. Examples

Below we demonstrate the power of the log derivative approach by considering two examples commonly found in biocontrol literature. We show stability properties could often be obtained at a glance. Homeostasis and disturbance rejection properties of such circuits are left to the next section.

#### 1) Incoherent feedforward circuit

Incoherent feedforward (IFF) circuits are both widely found in natural circuits [27], [28] and commonly used in synthetic circuit designs [29]–[31] to achieve homeostasis. We consider a simple IFF circuit below called the sniffer model [28]:

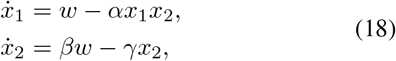

where *x*_2_ catalyzes degradation of *x*_1_, and *w* is a disturbance to the circuit. The disturbance *w* is considered a constant parameter for stability analysis.

To analyze this circuit’s structural stability, we first write down its log derivative matrix using the basic calculation rules established in Sectio II-C:

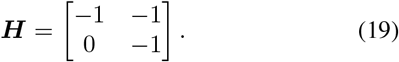

Note that this is a simple birth-death system, with constant log derivatives. Immediately, from Corollary 6, as ***H*** is triangular, we conclude that any positive fixed point of this circuit is structurally stable.

We emphasize the significance of this observation. Simply by a glance, we have determined that any positive fixed point of the system in Eq (18) is stable for the parameters *α, β, γ* and *w* taking arbitrary positive values. Furthermore, the functional forms of the production and degradation rates in (18) could vary as long as the log dervatives matrix is kept the same.

For example, that structural stability is robust to changes in functional forms could be used to find alternative implementations of the same structure. A variant of the IFF motif is the following (adapted from [31]):

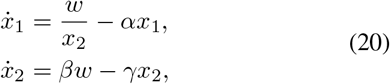

where *x*_2_ inhibits the production of *x*_1_ instead of degrading it. We see that the log derivative matrix for this system is the same as ***H*** in Eq (19), therefore sharing the same fixed point stability properties. Note how the structural stability property is robust to changes in the functional form of the production and degradation rates of *x*_1_ here. In Section IV-D we show that their homeostasis properties are the same as well.

Above examples of IFF circuits are both simple birth-death systems, where production and degradation rates have constant log derivatives. When attempting to write more realistic models, Hill functions that capture saturations are often used. For example, the *x*_1_ dynamics in Eq (20) could be modified as

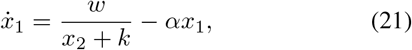

where *k* signifies the threshold for saturation of *x*_2_’s inhibition of *x*_1_. *k* can be physically interpreted as the binding strength between molecules *X*_2_ and the enzyme that catalyze the production of *X*_1_, which may correspond to *w*.

From Section II-C, we know the log derivative of production rate of *x*_1_ then satisfies 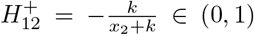, therefore the log derivative at a positive fixed point is

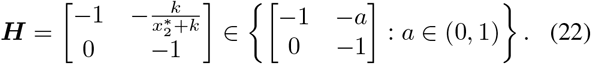

Although the exact value of ***H*** depends on 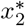 and *k*, since all ***H*** in the set in Eq (22) satisfies Corrolary 6, we still can conclude that any positive fixed point i s s tructurally stable.

#### 2) Sequestration negative feedback circuit

[32] proposed a circuit design based on molecular sequestration that could serve as a control module achieving homeostasis for general classes of plants connected to it. This architecture and its tradeoffs is analyzed in [33], [34], and [35] successfully implemented it in bacterial cells. We consider a simple example below containing the sequestration controller to illustrate how structural stability could be used to to analyze this control module with generic plants.

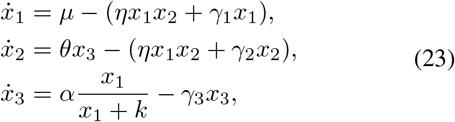

where *x*_3_ is to be controlled, and *x*_1_ and *x*_2_ sequester each other into a complex to be degraded. *x*_2_ senses *x*_3_’s concentration as *x*_3_ catalyzes the production of *x*_2_, while *x*_1_ actuates *x*_3_ to track reference *μ* by catalyzing the production of *x*_3_. Here we added self-degradation or dilution of *x*_1_ and *x*_2_ and used a Hill function for the production rate of *x*_3_ to make the model more realistic.

Using facts from Section II-C, we directly write that the log derivative at a positive fixed point satisfies

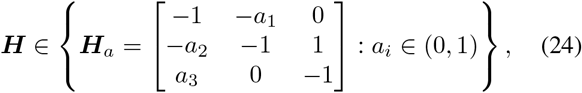

where *i* = 1, 2, 3. The particular values of *a*_*i*_ depend on the fixed point and the parameters, e.g.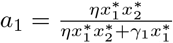. Scanning through values of *a*_*i*_ to find ***H***_a_ that is structurally stable could quickly characterize physical conditions that yield structural stability, providing guidelines for circuit design and implementation. For example, [32] argued that homeostasis is guaranteed by the circuit when *a*_1_ = *a*_2_ = 1, which corresponds to no dilution or degradation of the controller molecules *X*_1_ and *X*_2_. However, under this condition, ***H***_a_ is not diagonally stable for any values of *a*_3_, as tested from numerical computations. In contrast, if *a*_1_*, a*_2_ < 1, such as 0.99, then Re***H***_a_ could easily become negative definite for a wide range of *a*_3_ around 1. This simple computation demonstrates that dilution of the controller molecules, albeit damaging to the disturbance rejection property of the controller, significantly improves stability of the closed loop system. This enhances the observations in [33], [34].

## IV. Control through Structure

We have shown that fixed points’ stability could be quickly determined from structure. In the following, we show that robust perfect adaptation (RPA), biologists’ term for step disturbance rejection, can be implemented through structures.

### A. Birth-death control systems

We start by extending closed birh-death dynamical systems to open birth-death control systems.

#### Definition 8

A *birth-death control system* is

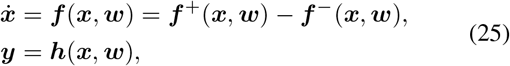

where 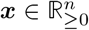 is state, 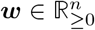 is disturbance input, and 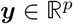 is output. The analytic functions 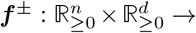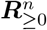 are production and degradation rates, and 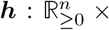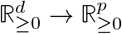 is output function.

Note that to keep the biological interpretation of the disturbance and output as coming from rates and concentrations, the variables ***w*** and ***y*** are also assumed to be non-negative.

In the following, we restrict our attention to the single-intput-single-output (SISO) case, so *w* and *y* are scalars.

### B. Linearized dynamics in terms of log derivatives

We assume the birth-death control system admits a positive reference point 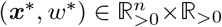 such that ***f*** (***x***^∗^*, w*^∗^) = 0 with positive outputs and rates: ***f*** ^±^(***x***^∗^*, w*^∗^) *>* 0, *y*^∗^ := *h*(***x***^∗^*, w*^∗^) *>* 0. Linearizing the system at this reference point then yields the following system.

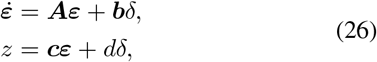

where 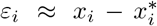, *δ* ≈ *w* − *w*^*^, *z* ≈ *y* − *y*^∗^ are linearized variables of *x*_*i*_, *w* and *y*. The matrices are ***A*** = *∂*_***x***_***f***(***x***\*, *w*\*) ∈ ℝ^*n* × *n*^, ***b*** = *∂*_*w*_***f***(***x***\*, *w*\*) ∈ ℝ^*n* × *1*^,***c*** = *∂*_*x*_*h*(***x***\*, *w*\*) ∈ ℝ^*n* × *1*^, and *d* = *∂_w_h*(***x***^∗^*, w*^∗^).

To express the linearized dynamics in log derivatives, we change variables into fold-change instead of additive difference:

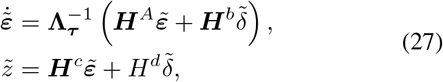

where 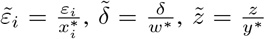 are fold-change linearized variables of *X*_*i*_, *w* and *y*. The log derivative matrices are

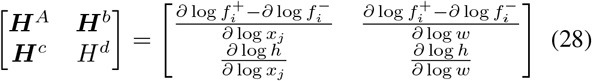

with the right hand side functions evaluated at (***x***^∗^*, w*^∗^). 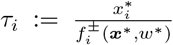 is the reference time-scale as before. In particular, the following relates the two versions of linearized dynamics:

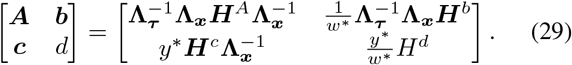

From this, we see that the transfer function for the fold-change linearized system is

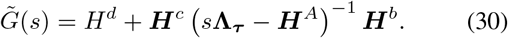

In fact, the fold-change transfer function is proportional to the traditional transfer function of the additive variables: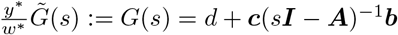

These calculations prepare us to discuss robust perfect adaptation based on structure.

### C. Structural robust perfect adaptation

Maintaining homeostasis despite uncertainties and disturbances is an essential function for biological organisms. Because of this, biomolecular circuits that achieve robust perfect adaptation (RPA) have been actively studied in systems biology [27]–[29], [36] and synthetic biology [30], [31]. In particular, one implementation of RPA through molecular sequestration has been proposed and implemented successfully in bacterial cells [32], [35], signifying important progress in principled design of biomolecular circuits.

On the other hand, our theoretical understanding of RPA in biomolecular systems is far from complete. From established tools of control theory, RPA as step disturbance rejection is thoroughly understood for linear dynamical systems, and the internal model principle could be used as a guideline for nonlinear systems [37]. However, biological constraints on implementable dynamics, such as variables need to be positive, make the design and implementation of RPA biomolecular circuits a challenging problem in general [30], [35]. Although significant progress has been made in RPA design from nonlinear analysis of biomolecular circuits, such approaches sensitively rely on the functional form of the production and degradation rates assumed in the model. More fundamentally, most biomolecular circuits are known to have desired properties like RPA only under certain parameter and state conditions, yet RPA based on the internal model principle requires RPA to hold globally. In comparison, RPA for linearized dynamics is local in nature, and it is described in a way that is independent of the rates’ functional forms. Below, we show how RPA in linearized dynamics could be robustified by implementing it through structure.

RPA, i.e. step disturbance rejection, in a linear system corresponds to the transfer function evaluating to zero at zero frequency. This means *G*(0) = *d **cA***^−1^***b*** = 0, which could be written as the determinant of a matrix, therefore the following definition and proposition.

#### Definition 9

A birth-death control system has *RPA* at reference point (***x***^∗^*, w*^∗^) if its linearized dynamics at this reference point in Eq (26) satisfies *z*(*t*) ⟶ 0 as *t* ⟶ ∞ for all constant disturbances *δ* ∈ ℝ.

#### Proposition 10 ([18], [30], [36])

A birth death control system has RPA at reference point (***x***^∗^*, w*^∗^) if and only if

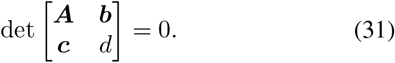

With Eq (27) in mind, we see log derivatives again nicely isolate the dependence on rates into the time-scales ***τ***, prompting the following definition of structural RPA.

#### Definition 11

A birth-death control system has *structural RPA* at reference point (***x***^∗^*, w*^∗^) if it is RPA at this point for all non-negative analytic rate functions ***f*** ^±^ and *h* that keep the log derivatives (***H***^A^, ***H***^b^, ***H***^c^, ***H***^d^) invariant at (***x***^∗^*, w*^∗^).

Practically, the variations and uncertainties described in the above definition come down to the time scales ***τ*** taking all positive values.

#### Theorem 12

A birth-death control system has structurral RPA at reference point (***x***^∗^*, w*^∗^) if and only if

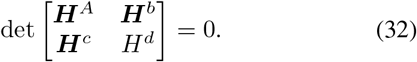

*Proof:* It is necessary that the fold-change transfer function satisfies 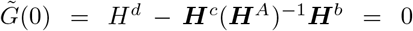, which is equivalent to Eq (32). Then since this condition is independent of ***τ*** and only depends on the log derivatives matrice, we see it is also sufficient for structural RPA.

### D. Examples

We continue the IFF and sequestration examples discussed in Section III-B. Let us consider the IFF system in Eq (18) with output *y* = *x*_2_, which states that we desire *x*_2_’s steady state concentration to be independent of disturbance *w*. Then the log derivative matrices satisfy

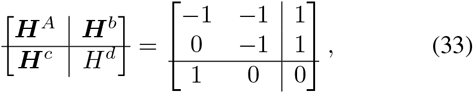

where horizontal and vertical rules are added for clarity. We calculate the determinant to be zero, concluding that the circuit is structurally RPA.

We emphasize the significance of this statement. After a glance at the system, we could write down the log derivatives and calculate the determinant, immediately concluding that *x*_1_’s steady state concentration is independent of step disturbances *w*, for the parameters *α, β, γ* taking arbitrary positive values. A simple calculation of log derivatives is able to provide powerful conclusions.

Structural RPA can be robust to variations in the functional forms of the model as well. This can used to suggest alternative implementations of the same RPA structure. Considering the variant in Eq (20), we see that the log derivative matrices are the same regarding the disturbance *w* and output *y* = *x*_1_, therefore having the same structural RPA property.

Furthermore, the structural mindset can help us understand how properties like RPA are valid under one regime while invalid in other regimes. Consider the variant in Eq (21) that captures saturation through a Hill function. When 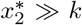, the log derivative matrices in Eq (33) hold asymptotically. So the regime with parameter condition 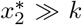 could be considered as the structual RPA regime. For any variations in parameters and functional forms of the system, as long as the system is still inside this regime, the structural RPA property holds. This suggests the view that a general birth-death system consists of several regimes, each with properties such as RPA implemented in its structure. This view is further explained and illustrated in [15].

Lastly, for the sequestration feedback circuit in Eq (23), since the goal is to make the concentration of *x*_3_ asymptotically track *μ*, we choose the output to be 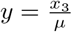. This yields log derivative matrices

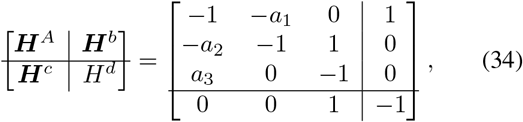

where *a*_*i*_ ∈ (0, 1) for *i* = 1, 2, 3. As argued in [32], making *a*_1_ = *a*_2_ = 1 achieves RPA. Indeed, the determinant of the above matrix is 1 − *a*_1_*a*_2_ − *a*_3_(1 − *a*_1_). When *a*_1_ = *a*_2_ = 1, this is zero. Therefore, *a*_1_ = *a*_2_ = 1 achieves structural RPA that is independent of *a*_3_ and all variations in parameters and rates’ functional forms, as long as the log derivatives are kept invariant.

## V. Discussion

In this work, we argued that structures of biomolecular circuits could be captured through log derivatives. We also demonstrated that fixed point stability and step disturbance rejection can be analyzed and designed through structure, independent of large variations in parameters and functional forms of circuit models.

This work builds on a train of thought that can be traced back to the very beginning of systems biology. Michaelis-Menten showed that time-scale separation and large concentration differences reveal distinct operating regimes of enzymatic regulations [8], [9]. In his pioneering work at the early days of systems biology [38], Savageau argued for the use of log dervatives as sensitivities of steady state concentrations to parameters, in order to study robustness. Furthermore, Savageau championed the view that a complex biomolecular circuit consists of several operating regimes. In particular, the concept of power systems is proposed that directly motivated the definition of birth-death systems and simple birth-death systems in this work. These pioneering ideas were later continued in gene regulation networks by Alon [29], and in stochasticity by Paulsson [24]. Works on these fronts became the foundational concepts and tools for systems biology.

This work, as well as several ongoing works, are attempts at formalizing many of the inspiring ideas from this train of thought, borrowing and creating tools from control theory, chemical reactions networks, and mathematics in the process. This formalization process also reveals further implications and connections. For example, Section II-B argues that log derivatives have their meaning rooted in the structures of biomolecular circuits, which can be formalized through time-scale separation with the application of implicit function theorem. This not only reveals the fascinating observation that log derivatives might form polytopes in general, but also provide a formal connection between powers / exponents, vertices of log derivative polytopes, and operating regimes of biomolecular circuits. Section III and IV demonstrate in a formal way that the structures captured by log dervatives have strong robustness properties to uncertainties in rates and functional forms used in the model, significantly extending the scope of the structural view for analysis and design of biomolecular circuits.

Another train of thought that this work borrowed much from is the theory of chemical reaction networks (CRNs) [16], [39]. With strong mathematical rigor, many fascinating developments recently appeared from this community on robustness and stability based on graphical structures of CRNs [21], [40]–[42]. One challenge of CRN theory is to identify a class of biological CRNs to avoid pathologies from extreme-case CRNs [41]. This work suggests that a candidate for a biological subset of CRNs could be the set of binding and catalysis reactions. From the analysis in Section II-B, a time-scale separation argument connects the biological structures underlying systems biology models to the graphical structures of CRN theories.

Lastly, there are many exciting questions left answered. Although the idea that regimes and simple birth-death systems could be viewed as approximations of the full system is introduced here through heuristic arguments, it could be rigorously developed using the framework of dissipative control [43]. Also, a foundational question is what classes of chemical reaction networks admit polytopic log derivatives. These questions are worth investigating, and we hope to answer them in another occasion in the near future.

## Acknowledgement

The authors would like to thank Daniele Cappelletti and John Marken for constructive discussions. F. X. and J. C. D. are partially funded by the Defense Advanced Research Projects Agency (Agreement HR0011-16-2-0049 and HR0011-17-2-0008).

